# Complete genome sequence of *Oryctes rhinoceros* Nudivirus isolated from Coconut Rhinoceros Beetle in the Solomon Islands

**DOI:** 10.1101/834036

**Authors:** Kayvan Etebari, Igor Filipović, Gordana Rašić, Gregor J. Devine, Helen Tsatsia, Michael J. Furlong

## Abstract

*Oryctes* Nudivirus (OrNV) has been an effective biocontrol agent against the insect pest *Oryctes rhinoceros* (Coleoptera: Scarabaeidae) for decades, but there is evidence that resistance could be evolving in some host populations. We detected OrNv infection in *O. rhinoceros* from the Solomon Islands and used Oxford Nanopore Technologies (ONT) long-read sequencing to determine the full length of the virus genomic sequence isolated from an individual belonging to a mitochondrial lineage (CRB-G) that was previously reported as resistant to OrNV. The complete circular genome of the virus consisted of 125,917 nucleotides, encoding 130 open reading frames (ORFs) for proteins in the range ~5 kDa (51aa) to ~140 kDa (1280aa). This Solomon Islands isolate has high (99.85%) nucleotide sequence identity to the previously sequenced isolate PV505 from Southern Luzon, Philippines, and has 138 amino acid modifications when compared with the originally described full genome sequence of the Ma07 strain from Malaysia.

---

The Nudiviridae form a monophyletic but highly diverse group of unassigned, large circular double-stranded DNA viruses, with enveloped and rod-shaped virions that are pathogenic for a wide range of arthropods (Burand, 1998; Wang et al., 2012). Currently, very few insect nudiviruses have been described, but the coconut rhinoceros beetle virus (*Oryctes rhinoceros* NV, OrNV) has been used as a pest biocontrol agent for decades (Bedford, 1986). This virus, previously known as *Oryctes* baculovirus, was isolated from Malaysian populations of *O. rhinoceros*, in the early 1960s and was successfully introduced into *O. rhinoceros* populations across the South Pacific islands in order to reduce pest damage to coconut palms (Bedford, 2013; Huger, 2005; Marschal, 1970). OrNV infection can be fatal to *O. rhinoceros* larvae, pupae and adults, but the disease is chronic and external symptoms in adult insects can be obscure (Burand, 1998).

The geographic range of the coconut rhinoceros beetle, *O. rhinoceros*, in the Pacific has recently expanded beyond its previous distribution (Fiji, Papua New Guinea, Samoa and Tonga) and now includes Guam (2007), Hawaii (~2013), Solomon Islands (2015), and more recently Vanuatu (2019) and New Caledonia (2019). It has been suggested that this resurgence is related to an *O. rhinoceros* mitochondrial lineage, termed the CRB-G haplotype, that is considered resistant to OrNV(Marshall et al., 2017; Reil et al., 2018). To obtain a better insight into the geographic diversity in OrNV, we sequenced the genomic DNA material of a virus-infected *O. rhinoceros* adult beetle collected in Solomon Islands.

A female adult beetle was collected from Guadalcanal, Solomon Islands using a pheromone trap (Oryctalure, P046-Lure, ChemTica Internacional, S. A., Heredia Costa Rica) in January 2019 and preserved in 95% ethanol. Total DNA was extracted from mid-gut tissue using the Qiagen Blood and Tissue DNA extraction kit, and was used for OrNV detection and beetle mitochondrial DNA amplification and sequencing. The *O. rhinoceros* mitochondrial haplotype was established as CRB-G (Marshall et al., 2017) *via* Sanger sequencing of the partial COXI gene sequence that was amplified using the universal barcode primers LCO1490 and HCO2198 (Folmer et al., 1994). Presence of the *Oryctes* Nudivirus (OrNV) was confirmed by successful amplification of a 945 bp product using the OrV15 primers that target OrNV-gp083 gene (Richards et al., 1999). The PCR product was visualized by gel electrophoresis and validated by Sanger sequencing. For ultra-long read DNA sequencing, we extracted high-molecular weight DNA from four legs and partial thorax tissue using a magnetic (SPRI) bead-based protocol. Specifically, smaller pieces of tissue (50 mm^3^) were each incubated in a 1.7 ml eppendorf tube with 360 μL ATL buffer, 40 μL of proteinase K (Qiagen Blood and Tissue DNA extraction kit) for 3h at RT, while rotating at 1 rpm (end-over-end). Then, 400 μL of AL buffer was added for 10 min, followed by 8 μL of RNase A for 5 minutes. Tissue debris was spun down quickly and 600 μL of homogenate was transferred to a fresh tube, where 600 μL of SPRI bead solution was added and incubated for 30 min while rotating at 1 rpm (end-over-end). After two washes with 75% ethanol, a final volume of 50 μL TE buffer was added. DNA quality was assessed on the 4200 Tapestation system (Agilent) and the concentration was determined using the Qubit broad-range DNA kit. DNA size selection (enrichment of DNA >10 kb) was done using the Circulomics Short Read Eliminator XS kit. Library preparation was done with 1 μg of size-selected HMW DNA, using the Ligation Sequencing Kit SQK-LSK109 (Oxford Nanopore Technologies, Cambridge UK) following manufacturer’s guidelines. In total, four libraries were loaded onto the MinION sequencing devices using the Flow Cell model R9.4.1 (Oxford Nanopore Technologies,) and sequenced by running the ONT MinKNOW Software. High-accuracy base calling on the raw sequence data was done with the Guppy base caller ONT v.3.2.4. High-quality sequences (=>Phred 13) were used for the genome assembly with Flye v.2.5 (metagenome assembly mode) (Kolmogorov et al., 2019).

Open reading frames in the resulting full-length circular consensus sequence were identified using CLC Genomics Workbench ver. 12.0 (QIAGEN) and then visually inspected and compared with the previously generated OrNV reference sequence (NC_011588). The NC_011588 reference genome sequence (named Ma07) was produced by multiple displacement amplification (MDA) of the whole viral genome from the homogenized mid-gut tissue of six infected insects collected from Johor, Peninsular Malaysia in 2007 (Wang et al., 2008). Finally, Nanopore long-read homopolymer stretches longer than 4 nucleotides were checked for artificial (sequencing technology-based) deletions or insertions and corrected based on the reference sequence.

Here, we produce 125,917 bp of the complete circular genome of *O. rhinoceros* Nudivirus using the Oxford Nanopore Technologies (ONT) long-read sequencing platform, with a median depth of 1,196 ×. The OrNV genomic sequence of the Solomon Island isolate has been deposited in GenBank under the accession number MN623374. ONT produced tens of single molecule reads ≥50 kbp that each span 40% or more of the entire virus genome (the longest read 98,6 kbp spanned 78% of the genome), ensuring unambiguous contiguity of the final assembly.

This new version of OrNV is 1,698 bp shorter than the genome previously reported by Wang *et al*., 2008. We identified 130 genes out 139 annotated coding regions in the genome of the Solomon Island isolate of OrNV. Seven previously annotated hypothetical proteins (OrNV_gp032, 68, 050, 081, 082, 066 and 091) did not produce any coding regions due to the presence of multiple stop codons and frame shifting in their new assembled sequence. Also, previously reported hypothetical proteins OrNV_gp129 and 130 are duplicated versions of OrNV_gp135 and 136 in our new assembly, suggesting MDA-related sequencing or assembly errors in the previous annotation that was prepared using shorter-read technology. We also found a rearrangement of four genes when compared to the previous assembly (Figure 1), supported by 446 single molecule reads (10□98.5 kbp long) that each span this entire region. This indicates that long read sequencing of non□amplified (raw) DNA extracts may be preferred when genomes of new DNA virus isolates are described in non□model organisms.

**Figure 1:**
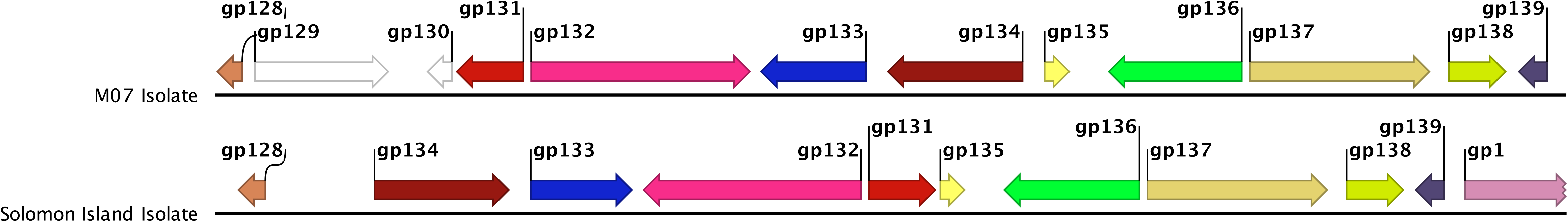
schematic diagram gene arrangement for Ma07 and Solomon Islands’ isolates. The SI isolate of OrNV is 1,698 bp shorter than Ma07. Two hypothetical proteins OrNV_gp129 and 130 are not present in new assembly-they were identified as duplicated versions of OrNV_gp135 and 136 in the Ma07 sequence.

In total, 352 single nucleotide polymorphisms (SNPs) have been found in 89 genes which caused 138 amino acid modifications in 53 coding regions in comparison with the previously reported OrNV sequence (Wang et al., 2008). Among these amino acid modifications, we found 37 amino acid deletions, 24 additions and 77 substitutions (Supplementary Table 1). 60% of identified genes in the newly sequenced virus have identical amino acid sequences with Ma07, while only 10% of genes have more than 4 amino acid modifications in their sequence. The greatest number of amino acid modifications have been found in OrNV_gp90 and OrNV_gp132, with 18 and 10 changes, respectively.

We also compared the new sequence to other available partial sequences from different OrNV isolates. We used the Jukes-Cantor algorithm to measure nucleotide distance under the UPGMA model and ran multiple gene alignment for regions of the Solomon Island isolate that overlapped with the complete genome sequences of the Malaysian isolate Ma07 (accession number NC_011588) and a partial genome sequence of The Philippines isolate PV505 (accession number AH015832) with only 40 open reading frames 37,051 bp, (Figure 3). Pairwise comparisons showed greatest similarity (99.85% nucleotide sequence identity) between the newly sequenced genome from Solomon Islands and PV505 (identity at 36,863 nucleotides, difference at 57 nt), while there were 84 nt differences out of 36,898 bp (99.77% identity) between PV505 and Ma07 (Figure 2 A-B). The phylogenic analysis based on partial sequence of DNA polymerase B, revealed significant similarity between the newly sequenced virus isolated in Solomon Islands and isolate PV505 (3 SNPs). Two other isolates from Sulteng and Riau, Indonesia (4 SNPs), and Ma07 (5 SNPs) are also closely related isolates (Figure 2 C).

**Figure 2:**
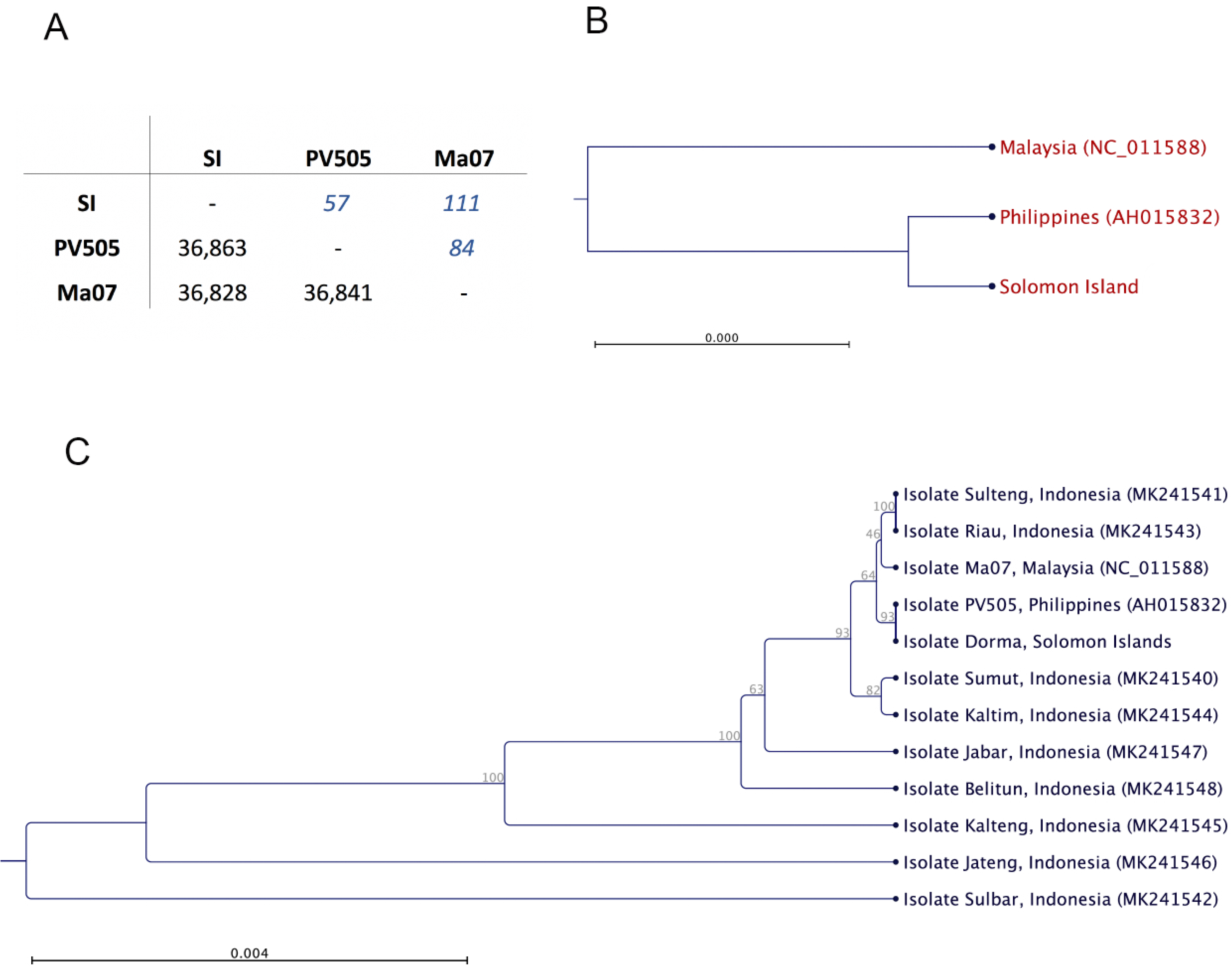
Phylogenic analysis and pairwise comparison of different geographical isolates of OrNV. **A)** Pairwise comparison of OrNV sequence of three geographical isolates (~37,000 bp overlapped fragment used for this alignment). Upper comparisons represent the number single nucleotide differences and lower comparisons are their level of identities. **B)** The cluster analysis shows more similarity between isolate sequenced from Solomon Island and Philippines. **C)** Phylogram produced based on partial sequence of DNA polymerase B.

**Figure 3:**
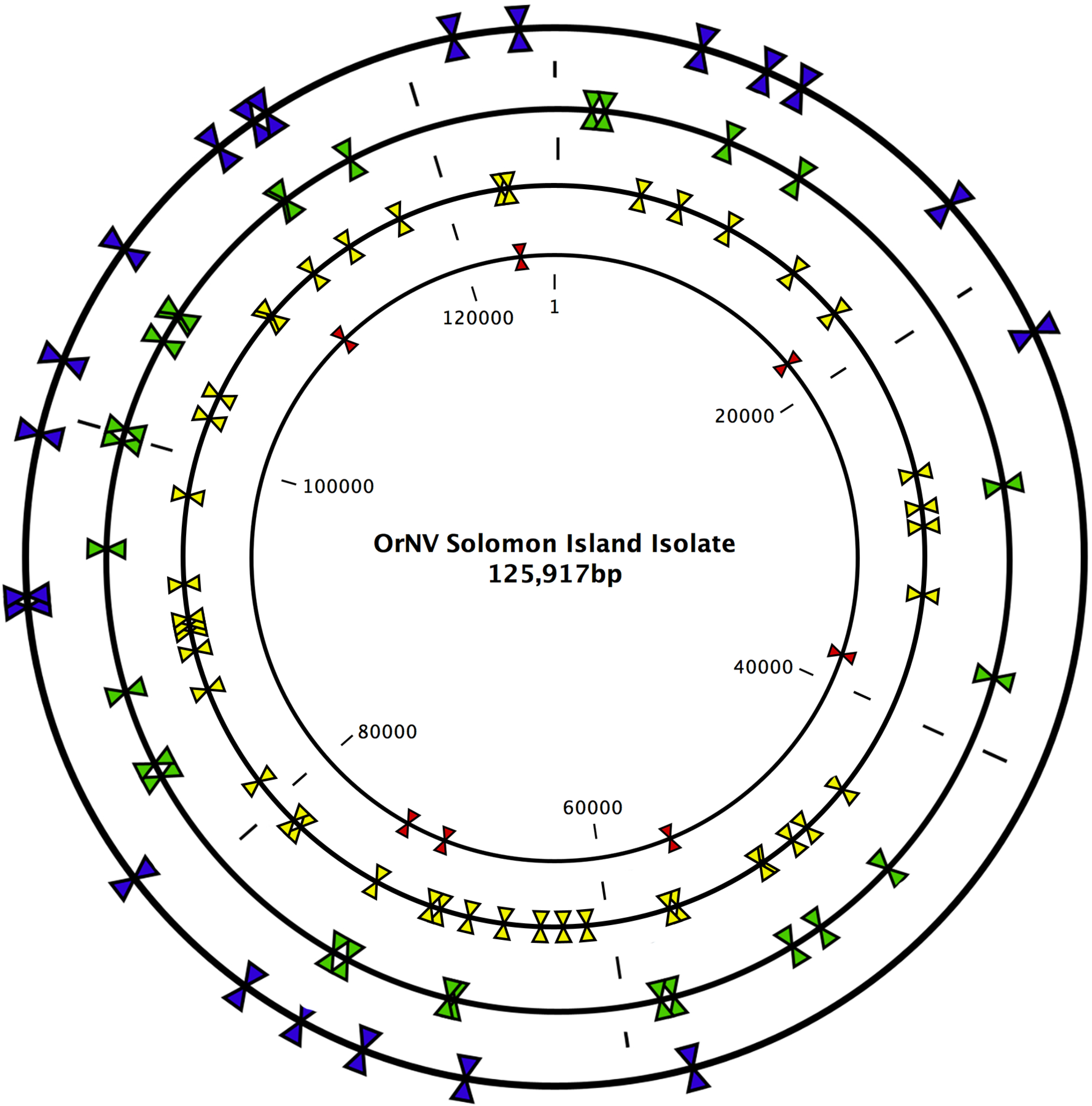
Restriction enzyme cleavage site map on the genomic sequence of OrNV isolate Solomon Islands. Blue, green, yellow and red color arrows represent the recognition sites for Bam HI, Hind III, Eco RI and Pst I, respectively.

Restriction fragment length polymorphisms have previously demonstrated the presence of genetic variation among field isolates of OrNV (Moslim et al., 2011). The *in silico* restriction endonuclease cleavage site map of the Solomon Islands OrNV isolate was produced with CLC genomic Workbench version 12 and revealed 43 EcoRI, 27 HindIII, 21 BamHI and 7 PstI fragments, respectively. In comparison, the previous version of the OrNV genome (Ma07) contains 44 EcoRI, 26 Hind III, 22 BamHI and 7 PstI cleavages sites (Wang et al., 2008).

Crawford and Zelazny (1990) did not find any changes in the *O. rhinoceros* viral genome < 2 years after the virus was introduced into in the beetle population in the Maldives. However, they reported some modifications, e.g. a point mutation and recombination over a longer, 4-year period (Crawford and Zelazny, 1990).

A study conducted on samples collected from Guadalcanal, Solomon Islands only three years prior to our collection did not detect OrNV *O. rhinoceros* (Marshall et al., 2017), suggesting very recent virus invasion into this host population. The source of the virus found in our 2019 collection is not known; it could have been introduced deliberately as a biological control agent or through accidental incursion of infected beetles from neighbouring islands. Interestingly, it was found in an individual from the CRB-G mitochondrial lineage, whose members have a low mortality rate from the OrNV infection (Marshall et al., 2017). Decreased mortality, however, is not necessarily related to host resistance to the virus, as many other factors could be involved in this phenomenon. During the course of evolution, viruses coevolve with their hosts to overcome host resistance and gain the upper hand in the evolutionary arms race (Hill and Unckless, 2018). Previously analyzed nudivirus genome data suggest that some genes, such as VLF-1, PIF-1, PIF-3, may be important in adapting to a new host (Hill and Unckless, 2018). While our isolate did not contain any changes in the amino acid sequences for PIF-1 or PIF-3 genes when compared to the Ma07 isolate, we did find five amino acid modifications (four addition and one deletion) in the VLF-1 gene (Supplementary table 1). Further sequence analyses of OrNV isolates in combination with the characterization of its effects on the host across multiple populations is needed to understand if such mutations compromise the efficacy of OrNV as a biocontrol agent that has kept the coconut rhinoceros beetle at bay across the Pacific for several decades. Our complete sequence and annotation of a newly found isolate in Solomon Islands will provide a valuable resource for such studies.

## Acknowledgements

This project was supported by the Australian Centre for International Agricultural Research funding (HORT/2016/185). and by core funds from the Mosquito Control Laboratory, QIMR Berghofer.

---

**Supplementary table 1: The list of identified coding regions on the genomic sequence of OrNV isolate Solomon Islands.** The nucleotide and amino acid modifications were compared with the originally described full genome sequence of the Ma07 strain from Malaysia.

